# Quantitative assessment of mesoscale cellular order and organization in the mouse hippocampus

**DOI:** 10.64898/2026.07.06.736467

**Authors:** Konstantin Hein, Humberto Romero-Limon, Conrad Möckel, Anne Karasinsky, Stephanie Möllmert, Jona Kayser, Alessio Zaccone, Jochen Guck, Tomohisa Toda

## Abstract

The hippocampus is characterized by a stereotypical macroscopic structure, where the nuclei are densely and heterogeneously packed among different subregions of the hippocampus. Despite the fact that tissue-specific cellular organization has been implicated in neural function, it has been technically challenging to quantitatively analyze mesoscopic cellular organization in the hippocampus due to its high cellular density. To overcome this technical hurdle, we developed Computational Biophysical Histomorphometry Software (CBHS), an automated image-analysis pipeline, aimed at quantifying nuclear shape and the order of the cellular ensemble in high-density areas. When applied to the subfields of hippocampus, we found that denser regions, most notably the dentate gyrus, were the most positionally, but least orientationally ordered. Nuclear shape exhibited a dependence on the local environment in a packing-dependent manner. This association was cell-type specific, with neurons, but not astrocytes displaying nuclear shape that varied with neighbour proximity, although astrocytes demonstrated greater intrinsic shape variance. The results reveal the presence of reproducible mesoscale cell packing order in hippocampal tissue, and are consistent with a nucleus-driven mechanical coupling between neighbouring cells. The present study provides a quantitative framework with which to understand mesoscopic tissue organization, thus enabling the formulation of testable hypotheses for future investigation.

## INTRODUCTION

Different brain areas show intrinsic differences in their overall topological properties. These properties emerge from the collective organization of individual cells. However, such organization remains sparsely quantified and rarely connected to the functional properties of the respective area. Recently, the functional significance of collective cellular organization in a brain area-dependent fashion has gained traction, including its potential role as a source of dysfunction in disease (Zhang et al., 2023; Das et al., 2023; Kim et al., 2024; Ferme et al., 2025). The hippocampus is one of the more investigated areas in terms of its cellular organization (Bayer et al., 1982; West et al., 1991; Keuker et al., 2003). It comprises three main subfields, Cornu Ammonis 1 (CA1), CA3, and the dentate gyrus (DG), that differ in function and cellular organization. At the macroscopic level, disease-induced topological alterations in the hippocampus are well documented, predominantly through tissue autopsy and functional magnetic resonance imaging (fMRI). Reduced functional connectivity, altered graph properties of the neural network, as well as changes in the density of hippocampal tissue have been linked to cognitive decline, memory deficits and neurological diseases such as Alzheimer’s disease (Allen et al., 2007; Wang et al., 2006; Tijms et al., 2013; Berron et al., 2021). Conversely, molecular disease mechanisms and targeted treatments are predominantly investigated at the microscopic scale, where individual cell morphology, subcellular structure and molecular environment are directly accessible. What remains poorly understood is how cellular-level changes connect to the macroscopic whole-tissue neuroimaging signatures. Fixed-tissue histological assessments offer a window to assess mesoscale changes between these two scales, thereby providing the resolution to quantify cellular ensembles while reflecting the aggregate micro-mesoscopic structural properties that shape macroscopic signatures. Bridging all three scales is essential for translating cellular and molecular findings into interpretable macroscopic signatures of disease.

At the mesoscale, cell packing and density have been shown to be valuable qualitative read-outs of broader changes in hippocampal structure across several disease models (Wishart et al., 2010; Baglietto-Vargas et al., 2010; Arranz et al., 2014; Schoene-Bake et al., 2014). It has thus been established that the significance of mesoscopic tissue organization is associated with brain dysfunction. However, this association is primarily established through qualitative or semiquantitative methodologies, without the establishment of a link to other scales. It is believed that mature post-mitotic neurons in the CNS do not migrate or move, outside of a few microns over prolonged time periods (Rakic, 1974). However, when many tightly packed neurons move by a few microns, the cumulative effect on the spatial organization of the collective tissue can be significant. For example, stereological studies have shown that the number of granule cells and their packing density in the DG increase from juvenile to adult life, and that age-related changes in subfield volume and neuron number can be partially decoupled (Bayer et al., 1982; West et al., 1991; Keuker et al., 2003). These results indicate that cellular spatial organization might be more dynamically altered than previously assumed, not following a fixed trajectory, but rather reacting to external circumstances, such as experiences and disease.

To investigate such a dynamic system at the mesoscale, different metrics can be used, which all relate to the distance between cells, their angles and their shape. Metrics, such as the radial distribution function or local nematic order, enable the quantification of the spatial cellular organization in tissue. Furthermore, they can provide insights into the relationship between the spatial organization and the density of cells in the area of interest. Intriguingly, hippocampal neurons exhibit very high nuclear volume fractions within their cell bodies (Bayer et al., 1982; West et al., 1991; Keuker et al., 2003). Nuclei in these areas are also known to dynamically change their morphology based on disease and functional states (Das et al., 2023). Tissues with similarly high nuclear volume fraction and similarly tight packing, such as the zebrafish retina, have demonstrated a nucleus-driven tissue organization, including a relationship between cellular packing order and density (Kim et al., 2024; Ferme et al., 2025). Furthermore, these studies demonstrated that the cellular spatial organization itself might be functionally relevant and dynamically altered. This might imply that other neuronal cells outside the retina might also order themselves beyond a certain density threshold in functionally relevant manner. Together, it is conceivable that quantifying changes in the cellular packing order and density of cells in the hippocampus represents the first steps towards understanding the spatial organization of nuclei. By connecting different scales, we would gain insight into the functional relevance of the mesoscale itself and therefore form a bridge between the molecular and tissue-level changes. However, a unified quantitative framework for the mesoscale analysis of hippocampal structure is currently lacking.

To address this technical and knowledge gap and to investigate the relationship between cellular packing order and density in the highly-packed hippocampal tissue, we developed the Computational Biophysical Histomorphometry Software (CBHS). CBHS is an all-in-one pipeline for the analysis of confocal images, integrating background correction, CellPose-based nuclear segmentation, 3D reconstruction, fluorescence intensity correction and automated marker-positive cell filtering. It also provides a unified framework for quantifying the nuclear shape, density, and cellular packing order of nuclei across hippocampal subfields (Pachitariu et al., 2025; Kleinberg et al., 2022; Weigert et al., 2020). Using CBHS, we characterized the mesoscale organization of the hippocampus quantitatively and found that, similar to the zebrafish retina, cells in the mouse hippocampus, show an order in their cellular organization beyond a theoretical density threshold. Furthermore, the shape of the nuclei changes under the influence of the local environmental structure. Ultimately, this provides a basis for analysing dynamic and potentially functionally relevant changes in the cellular organization of these neuronal ensembles. These findings revealed the principles for investigating the trajectories of brain subregion-specific cellular and tissue organization, which could be critical for tissue-specific function and plasticity.

## Material and Methods

### Mice handling, tissue processing and immunohistochemistry

All procedures relating to mouse care and treatment were approved by the Government of Saxony and performed in accordance with their guidelines. Three male C57BL/6J mice aged 4 months were used. Mice were euthanised with a lethal dose of sodium pentobarbital (WDT). After death had been confirmed, the animals were transcardially perfused with ice-cold 1× PBS, followed by 50 mL of freshly prepared 4% paraformaldehyde (PFA; Sigma-Aldrich) in PBS. Following perfusion, brains were post-fixed in 4% PFA overnight at 4 °C and then washed three times in PBS. Brains were transferred through a graded sucrose series in 0.1 M phosphate buffer for 24 hours for each step (15% followed by 30%); once the brains had sunk in the 30% solution, 40 µm coronal sections were cut on a sliding microtome (Leica SM 2010 R) cooled with dry ice. Sections were stored in cryoprotective solution (25% ethylene glycol (Roth), 25% glycerol (VWR) in 0.1 M phosphate buffer, pH 7.4) at -20°C until use. For immunostaining, free-floating sections were first washed twice in PBS on a plate shaker (15 min each). Antigen retrieval was then performed by transferring the sections to 10 mM citrate buffer (pH 6.0) and heating them in a vegetable steamer at approx. 90°C for 8 min, followed by a 10 min cool-down within the steamer. After two 5 min washes in PBS, sections were permeabilized in two 15 min rounds of 0.5% Triton X-100 in 1× PBS (permeabilization buffer), followed by a single round of combined permeabilization and blocking in 0.5% Triton X-100 with 3% horse serum in 1× PBS (blocking buffer). Primary antibodies (Table 1) were subsequently applied in blocking buffer for 48–72 hours at 4°C. Sections were then washed three times in permeabilization buffer (15 min each) and incubated for a further 30 min in blocking buffer. Secondary antibodies and fluorescent dyes (Table 1) were applied in permeabilization buffer for 2 h at room temperature. Finally, sections were washed three times in permeabilization buffer and twice in PBS, and mounted with Mowiol under 1.5H coverslips.

**Table 1:** Primary and secondary antibodies and dyes used in this study. All tissue was mouse.

| Antibody | Host / isotope | Dilution | Distributor, Cat. no. |
| --- | --- | --- | --- |
| *Primary antibodies* |  |  |  |
| Sox2 | Rat, monoclonal IgG2a | 1:2000 | ThermoFisher, 14-9811-82 |
| NeuN | Mouse, monoclonal IgG1 | 1:250 | Millipore, MAB377 |
| Ctip2 | Rat | 1:200 | Abcam, ab18465 |
| GFAP | Goat, polyclonal | 1:1000 | Abcam, ab53554 |
| S100β | Rabbit, monoclonal | 1:200 | Abcam, ab52642 |
| *Secondary antibodies and dyes* |  |  |  |
| Anti-Rabbit Cy3 | Donkey IgG | 1:500 | Jackson IR, 711-165-152 |
| Anti-Rat Cy5 | Donkey IgG | 1:500 | Jackson IR, 712-175-153 |
| Anti-Mouse A488 | Donkey IgG | 1:500 | Jackson IR, 715-545-150 |
| Anti-Goat A488 | Donkey IgG | 1:500 | Dianova, 705-545-147 |
| DAPI | — | 1:2000 | ThermoFisher, 1387190 |

### Image acquisition

All images were acquired on one of four systems: a Nikon CSU-W1 spinning-disc confocal or a Zeiss SD 1 spinning-disc system, each using camera-based detection, or a Zeiss LSM 980 or Zeiss LSM 900 point-scanning confocal, each detecting through photomultiplier tubes (PMTs). In every case, imaging was performed with either a 20x air objective (0.8 NA) or a 40× air objective (0.95 NA), and excitation was provided by the 405, 488, 561 and 640 nm laser lines. Laser power and detector gain were adjusted so that signal intensities occupied no more than one-third of the available 16-bit intensity range. Pixel sizes ranged from 0.1135 µm to 0.35 µm and *z*-step intervals from 0.16 µm to 0.34 µm. In all cases, the entire 40 µm section was imaged, together with an additional approximately 10% beyond each surface. Airyscan acquisition and subsequent deconvolution were applied for the LSM-based imaging. The process of image-analysis is described in the results section and in Table 2.

**Table 2.** Image analysis metrics.

| Metric | Formula | Description |
| --- | --- | --- |
| Illumination correction |  |  |
| Opening | $\gamma_B(I) = \delta_B(\epsilon_B(I))$ | Dilation ($\delta_B$) after erosion ($\epsilon_B$) of an image, used to eliminate small objects from the background calculation. |
| Gaussian filter | $G_\sigma = \frac{1}{2\pi\sigma^2} e^{-(x^2+y^2)/(2\sigma^2)}$ | Calculation of the illumination function for an image with a Gaussian blur scale $\sigma$. |
| Size of the structuring element | $q = \max\left(31, 2 \left\lfloor \frac{\text{base} \times \text{kernel fraction}}{2} \right\rfloor + 1\right)$ | Automatic calculation of the (odd) kernel size for the ellipsoidal structuring element $B$, ensuring a size of at least 31 pixels for smoothing and a user-defined kernel fraction. |
| Illumination correction | $C = \max(I - (\gamma_B(I) * G_\sigma), 0)$ | Removal of the illumination variation (the opening smoothed by $G_\sigma$), with the result clipped at 0. |
| Root mean square of residuals | $\text{RMS}_{\text{residual}} = \sqrt{\frac{1}{N} \sum_{i=1}^N (z_i - \hat{z}(x_i, y_i))^2}$ $= \sqrt{\frac{1}{N} \sum_{i=1}^N (z_i - (ax_i + by_i + c))^2}$ | Evaluation of background variation, where $N$ is the number of pixels, $(x_i, y_i)$ are the pixel coordinates and $z_i$ is the pixel value; $a, b, c$ are obtained via least squares, $(a, b, c) = \arg \min_{a,b,c} \sum_i (z_i - ax_i - by_i - c)^2$. |
| **2D Morphology** |  |  |
| Area | $A = N_{\text{px}} \cdot d_x d_y$ | Pixel count of the object multiplied by the physical pixel area. |
| Perimeter | $P = P_{\text{px}} \cdot \sqrt{d_x^2 + d_y^2}$ | Crofton-style perimeter from regionprops, scaled by the diagonal pixel length. |
| Major and minor axes | $M = \lambda_1 \cdot \sqrt{d_x^2 + d_y^2},$<br>$m = \lambda_2 \cdot \sqrt{d_x^2 + d_y^2}$ | The region is fitted to an ellipse using the second central moment $\mu_{pq} = \sum_{(x,y) \in R_k} (x - \bar{x})^p (y - \bar{y})^q$, with variance matrix $\Sigma = \begin{bmatrix} \mu_{20}/A & \mu_{11}/A \\ \mu_{11}/A & \mu_{02}/A \end{bmatrix}$, from which the eigenvalues $\lambda_1 > \lambda_2$ give the major and minor axes. |
| Centroid | $(x_c, y_c) = (\bar{x}_{\text{ROI}} + x_{\text{min}}, \bar{y}_{\text{ROI}} + y_{\text{min}})$ | Centre of mass of the object, mapped from the ROI back to full-image coordinates. |
| Orientation | $\theta = \frac{1}{2} \text{atan2}(2\mu_{11}, \mu_{20} - \mu_{02})$ | Angle (radians, $\theta \in [-\pi/2, \pi/2]$) between the row axis and the major axis of the fitted ellipse, i.e. the eigenvector direction of $\lambda_1$. |
| Circularity | $\text{CI} = \frac{4\pi A}{P^2}$ | Wadell-style shape factor; equals 1 for a perfect circle and decreases with roughness. |
| Aspect ratio | $\text{AR} = \frac{m}{M}$ | Ratio of minor to major axis; equals 1 for a circle, $\rightarrow 0$ for elongated shapes. |
| Aspect-ratio circularity | $C_{\text{AR}} = -0.528 \text{AR}^2 + 1.259 \text{AR} + 0.270$ | Empirical fit (Zheng & Hryciw 2015) giving the circularity expected from aspect ratio alone. |
| Roundness (corrected) | $R = \text{CI} + (0.913 - C_{\text{AR}})$ | Zheng & Hryciw corrected roundness that removes the aspect-ratio contribution from circularity. |
| Solidity | $s = \frac{A}{A_{\text{convex}}}$ | Ratio of object area to convex-hull area; quantifies concavity. |
| Elongation | $E = \sqrt{1 - \frac{m^2}{M^2}}$ | Eccentricity of the fitted ellipse (`regionprops.eccentricity`); 0 for a circle, $\rightarrow 1$ for a line. |
| Eccentricity (bbox) | $\varepsilon = \frac{\min(\Delta x, \Delta y)}{\max(\Delta x, \Delta y)}$ | Aspect ratio of the axis-aligned bounding box of the object. |
| Convex perimeter | $P_{\text{convex}} = \sum_{k=1}^{N_H} \ v_k - v_{k-1}\ \cdot \sqrt{d_x^2 + d_y^2}$ | Perimeter of the 2D convex hull $H(R_k)$ (QHull), summed as Euclidean edge lengths between successive hull vertices $v_k$ and scaled to physical units. |
| Convex area | $A_{\text{convex}} = \vert \{(x, y) \in \mathbb{Z}^2 : (x, y) \in H(R_k)\}\vert \cdot d_x d_y$ | Area of the 2D convex hull: number of integer lattice points contained in $H(R_k)$, scaled by the physical pixel area. |
| Convexity | $\text{Conv} = \frac{P_{\text{convex}}}{P}$ | Ratio of convex perimeter to actual perimeter; close to 1 for smooth objects, $< 1$ for rough ones. |
| Sphericity | $\Phi_{2D} = \frac{r_{\text{ins}}}{r_{\text{circ}}}$ | Ratio of inscribing to circumscribing radius, both measured from the centroid to the outline. |
| Mean curvature | $\bar{\kappa} = \frac{1}{N} \sum_{k=1}^N \ v_k - v_{k-1}\ $ | Mean edge length between successive convex-hull vertices; a coarse curvature proxy. |
| Bending energy | $E_{\text{bend}} = \sum_{k=1}^N \ v_k - v_{k-1}\ ^2$ | Sum of squared edge lengths between successive convex-hull vertices. |
| Number of neighbours | $N_{\text{neigh}} = \vert \{\text{masks}[p] : p \in \partial_i \pm k\hat{x}, \pm k\hat{y}\} \setminus \{0, i\}\vert $ | Count of unique non-self, non-background labels within $k$ px outside the object's outline along the cardinal directions. |
| **3D Morphology** |  |  |
| Centroid | $x_c = x_{\min} + \frac{\sum_{x,y} P_{xy}(x,y) x}{\sum_{x,y} P_{xy}(x,y)}$ $y_c = y_{\min} + \frac{\sum_{x,y} P_{xy}(x,y) y}{\sum_{x,y} P_{xy}(x,y)}$ $z_c = z_{\min} + \frac{\sum_{x,z} P_{xz}(x,z) z}{\sum_{x,z} P_{xz}(x,z)}$ | Centre of mass mapped from the ROI back to full-image coordinates, with $P_{xy}$ the $z$-projection and $P_{xz}$ the $y$-projection of the object. |
| Voxel scaling | $V_{\text{voxel}} = d_x d_y d_z,$ $d\ell = \sqrt{d_x^2 + d_y^2 + d_z^2}$ | Physical volume of a single voxel and the voxel diagonal length. Volumetric quantities are scaled by $V_{\text{voxel}}$; surface/length quantities by the relevant face areas or $d\ell$. |
| Volume | $V = N_{\text{vox}} \cdot V_{\text{voxel}}$ | Number of voxels $N_{\text{vox}}$ occupied by the object, scaled to the physical voxel size. |
| Surface area | $S_{\text{area}} = (\sum O + 4N_z) d_y d_z + (\sum_{z=0} I + \sum_{z=z_{\max}} I) d_x d_y$ | Outline voxels $\sum O$ plus corners ($4N_z$ per $z$-plane), plus the bottom and top caps of the object. |
| Sphericity | $\Phi = \frac{\pi^{1/3} (6V)^{2/3}}{S_{\text{area}}}$ | Ratio scaled so that a perfect sphere gives $\Phi = 1$ and any other shape a lower value. |
| Radial measurements | $\bar{r} = \frac{1}{N} \sum_{k=1}^N \ p_k - c\ ,$ $r_{\text{in}} = \min_k \ p_k - c\ ,$ $r_{\text{circ}} = \max_k \ p_k - c\ $ | Mean, minimum and maximum distance from the centroid $c$ to each outline point $p_k$. |
| Deformation | $D = \frac{r_{\text{in}}}{r_{\text{circ}}}$ | Ratio between the smallest concavity and the largest convexity of the object. |
| Major and minor axes | $L_{\text{major}} = \ell_{\text{major}} d\ell,$<br>$L_{\text{minor}} = \ell_{\text{minor}} d\ell$ | Longest and shortest principal axis lengths (from the inertia ellipsoid via `regionprops`), in physical units. |
| Aspect ratio | $\text{AR} = \frac{\ell_{\text{minor}}}{\ell_{\text{major}}}$ | Elongation in $(0, 1]$. Values near 1 mean roughly isotropic; small values mean rod- or disc-like. |
| Convex surface area and volume | $S_{\text{convex}} = A_{\text{hull}} d_x d_z,$<br>$V_{\text{convex}} = V_{\text{hull}} V_{\text{voxel}}$ | Surface area and volume of the convex hull fitted to the object's outline points; the “shrink-wrapped” references against which concavity is measured. |
| Convexity | $\text{Convexity} = \frac{V}{V_{\text{convex}}}$ | Ratio of object volume to convex-hull volume, in $(0, 1]$. Values near 1 mean a convex, smooth body; lower values indicate concavities or protrusions. |
| **2D Object intensity** |  |  |
| For each channel $c$ the global background is $\text{bg}_c = \frac{1}{A_{bg}} \sum_{(x,y) \notin \text{masks}} I(x, y)$, where $A_{bg}$ is the area with no detected object. | | |
| Mean intensity (background divided) | $\bar{I}_k = \frac{\frac{1}{A} \sum_{(x,y) \in R_k} I(x, y)}{\text{bg}_c}$ | Mean intensity inside the object normalised by the global background of that channel. |
| Mean intensity (background subtracted) | $\bar{I}_k = \frac{1}{A} \sum_{(x,y) \in R_k} I(x, y) - \text{bg}_c$ | Background-subtracted mean intensity inside the object, per channel. |
| Edge intensity | $\bar{E}_{I_k} = \frac{1}{\vert O_k\vert } \sum_{(x,y) \in O_k} I(x, y) - \text{bg}_c$ | Background-subtracted mean intensity along the object's outline $O_k$. |
| Radial bin $b$ | $I_b^c = \frac{\text{mean}(\text{ch}_c[\text{layer}_b])}{\sum(\text{ch}_{c,\text{ROI}} - \text{bg}_c)}$ | Mean intensity within layer $b$ of the Euclidean distance transform, normalised by the total background-corrected ROI signal. Bins are `linspace(0, $D_{\text{max}}$, $n_{\text{bins}} + 1$)` on $D = \text{EDT}(\text{ROI})$. |
| **Order** |  |  |
| *Given centroids $c$, $p_1$, $p_2$ and vectors $v_j = p_j - c$:* |  |  |
| Pair angle | $\theta = \arccos\left(\frac{v_1 \cdot v_2}{\ v_1\ \ v_2\ }\right)$ | Angle subtended at a cell centroid by two of its neighbours. |
| Local nematic order (neighbour pairs) | $n_{o,k} = \cos(2(\theta_k - \theta_{k-1}))$ | Pairwise nematic alignment between successive neighbour angles; averaged per cell. |
| Mean nematic order | $\langle n_o \rangle = \frac{1}{K} \sum_{k=1}^K n_{o,k}$ | Average local nematic order over all neighbour pairs of a cell. |
| Angular dispersion | $\sigma_\theta = \text{std}\left(\frac{\theta}{\bar{\theta}}\right)$ | Standard deviation of neighbour angles after normalisation by their mean. |
| Radial dispersion | $\sigma_d = \text{std}\left(\frac{d}{\bar{d}}\right)$ | Standard deviation of neighbour distances after normalisation by their mean. |
| Min neighbour distance | $d_{\min} = \min(d_1, d_2) \cdot \sqrt{\bar{d}_x^2 + \bar{d}_y^2}$ | Smallest distance to the two nearest neighbours, scaled to physical units. |
| Mean touching distance | $\bar{d}_{\text{touch}} = \text{mean}(\text{dist}(c_i, c_{j \in \mathcal{N}_i})) \cdot \sqrt{\bar{d}_x^2 + \bar{d}_y^2}$ | Mean distance between the cell and its outline-touching neighbours $\mathcal{N}_i$; falls back to the two nearest neighbours if none touch. |
| Mean 3-NN distance | $\bar{d}_3 = \text{mean}(d_1, d_2, d_3) \cdot \sqrt{\bar{d}_x^2 + \bar{d}_y^2}$ | Mean distance to the three nearest neighbours, scaled to physical units. |
| Radial distribution function | $g(r) = \frac{1}{\rho N} \cdot \frac{\sum_{i \neq j} \delta(r - \vert r_i - r_j\vert )}{2\pi r \Delta r}$ | Histogram of pairwise centroid distances, normalised by the 2D ideal-gas expectation $2\pi r \Delta r \rho$. |
| Two-point (excess-entropy) integral | $S_2 = -\frac{\rho}{2} \int_0^R [g(r) \ln g(r) - g(r) + 1] 2\pi r dr$ | The integrand peaks where $g(r)$ departs most from ideal-gas behaviour; small values indicate a structureless fluid. |
| Bond-orientational order | $\Psi_n(j) = \frac{1}{z_j} \sum_{k \in N(j)} \exp(i n \theta_{jk})$ | Halperin & Nelson (1978) local complex order parameter at symmetry order $n$, for nucleus $j$ with Delaunay-neighbour set $N(j)$ and bond angles $\theta_{jk} = \text{atan2}(y_k - y_j, x_k - x_j)$. |
| Nematic order (orientations) | $c_2 = \langle \cos 2\theta \rangle, \quad s_2 = \langle \sin 2\theta \rangle,$<br>$S_{\text{orient}} = \sqrt{c_2^2 + s_2^2}$ | Standard 2D nematic order parameter measuring alignment of the nuclei's long-axis orientations. |

### Statistics

Statistical analyses were performed in Python using SciPy, statsmodels, NumPy, and scikit-bio (v0.7.2), with all tests two-sided at *α* = 0.05. Because measurements were derived from large populations of segmented nuclei or sliding windows rather than from the three animals, consistency across animals was verified once and all reported inferences were computed at the population level. Significance was established by resampling rather than from parametric distributional assumptions.

Significance was decided by a single resampling criterion applied throughout the group comparisons and correlations: each test was recomputed over many size-matched subsamples, and an effect was called significant when at least 80% of iterations reached *p <* 0.05, with results graded by the significant-iteration fraction as ≥0.80 (*), ≥0.90 (**), and ≥0.95 (***). Size-matching placed groups of very different sizes on a common footing by drawing size-matched subsamples of 90–100 observations.

Differences in continuous, non-normally distributed data across three or more independent groups were assessed with a bootstrapped Kruskal–Wallis omnibus test, followed by Mann–Whitne U pairwise comparisons. Each iteration subsampled every group without replacement to a common size of 90–100 observations; the Kruskal–Wallis test was repeated 2 × 10^4^–1 × 10^5^ times and each Mann–Whitney U pair 5 × 10^3^–1 × 10^4^ times, with the significant-iteration fraction and the mean and median *p* recorded.

Differences between two paired (matched) measurements of the same units were tested with the two-sided Wilcoxon signed-rank test, reporting the rank-biserial correlation as the effect size; a paired *t*-test was added as a parametric complement. The agreement between two paired measurements was summarized with the mean absolute error (MAE) and the root-mean-square error (RMSE).

The association between two continuous variables was quantified with Spearman’s rank correlation *ρ* for monotonic dependence, and with Pearson’s *r* where a linear relationship was of interest. The corresponding trend was fitted by ordinary least-squares regression (slope and *R*^2^) for the linear component and by locally weighted scatterplot smoothing (LOWESS; smoothing fraction 0.30) for non-linear dependence. The 95% bands for the LOWESS and linear fits, and the 95% intervals for the per-region slopes and correlation coefficients in the forest plots, were obtained by percentile bootstrap using 200 fixed resamples of 1500 points. Whether an association persisted after accounting for a covariate was tested by partial-residual analysis: the response was regressed against the predictor, and the per-group residuals were plotted against the predictor as a LOWESS curve with a 95% bootstrap band. Each group’s mean residual was tested against zero with a one-sample *t*-test, and the residual–predictor Spearman correlation was reported as a check for remaining structure.

Multivariate differences between groups in a high-dimensional feature space were examined by principal component analysis (PCA), computed by singular value decomposition of the *z*-scored feature matrix. The first two principal components were retained, and each group’s spread was depicted as a 2*σ* covariance ellipse, enclosing approximately 86% of the mass of a bivariate-normal distribution. Differences in distribution location were tested by permutational multivariate analysis of variance (PERMANOVA) on Euclidean distances with 999 label permutations, reporting the proportion of multivariate variance attributable to the grouping (*R*^2^) as the effect size. Differences in multivariate dispersion were tested separately by a permutational test of multivariate dispersion (PERMDISP), quantified as the mean distance of each point to its group centroid, with Levene’s test on a one-dimensional composite (computed on class-balanced sub-samples) as univariate corroboration. Both tests were performed in the plotted PC1–PC2 space and, as a sensitivity analysis, in the full *z*-scored feature space, with group sizes equalized by repeated balanced subsampling. The Euclidean PERMANOVA was evaluated through its closed-form sum-of-squares decomposition and verified against the scikit-bio (v0.7.2) reference implementations on balanced subsamples to numerical precision.

Effect sizes and coefficients — epsilon-squared (*ε*^2^), Cliff’s delta (*δ*), the rank-biserial correlation (*r*_rb_), Spearman’s *ρ*, the OLS slope, the segmented-regression breakpoint, and the mean difference 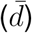— were reported with a bootstrap 95% confidence interval, whereas the median or mean of a measured quantity was reported with its central 68% spread (the 16th–84th-percentile interval). Both were written in the compact asymmetric form 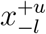, in which the upper and lower offsets are the distances from the central value to the bounds of the corresponding interval (for a 68% spread, *u* = *P*_84_ − *P*_50_ and *l* = *P*_50_ − *P*_16_; for an effect size, the bounds of its bootstrap 95% CI). Offsets were rounded to two significant figures, with the central value given to matching precision.

Although the window- and nucleus-level analyses are drawn from a single segmented image set, the reported sample size for any given panel reflects only those units for which the specific metric could be defined correctly. Consequently, two window-based panels can differ in N when one metric is undefined for a subset of windows. For example, mask-clipped nuclear density requires sufficient overlap with the annotated subfield mask and is therefore unavailable for partial-edge or low-occupancy windows, whereas the radial-distribution-derived positional (*S*_2_), coordination (*Z*), and orientational (*S*) measures are computed across the broader set of placed windows, yielding a larger denominator. The same issue produces differing N among nucleus-based panels, where each comparison retains only nuclei satisfying that panel’s inclusion criteria—namely complete shape-descriptor vectors for the principal-component analysis, marker-positive cell-type with or without the co-positive labelling for the cell-type comparisons, and restriction to a given density regime or hippocampal subfield.

## RESULTS

To investigate the mesoscale organization of tissues, it is necessary to be able to resolve individual cells/nuclei while simultaneously quantifying their spatial relationships. Hippocampal subfields present several inherent challenges for automated nuclear analysis. First, the uneven cellular distribution produces non-uniform background intensity and autofluorescence (i). Second, the tight packing of nuclei make it difficult to conduct instance segmentation, particularly where adjacent objects overlap (ii). Third, optical sectioning truncates the polar caps of nuclei in 3D because intensity in the surrounding plane exceeds that at the top and bottom of each body (iii), leading to systematic under-representation of nuclear volume. To address these limitations while enabling quantitative analysis of the spatial tissue organization across diverse cellular ensembles, we developed CBHS, a pipeline that combines per-slice background correction across the *Z* -stack, 2D segmentation of DAPI-stained nuclear images with a CellPose CPSAM model fine-tuned using hand-annotated nuclei in the DG, mask-guided edge enhancement of the raw DAPI volume prior to 3D reconstruction with Stardist 3D, and an in-house mask-cleaning and interpolation step termed Yukimatsu (Figure 1, for an overview).

**Figure 1:**
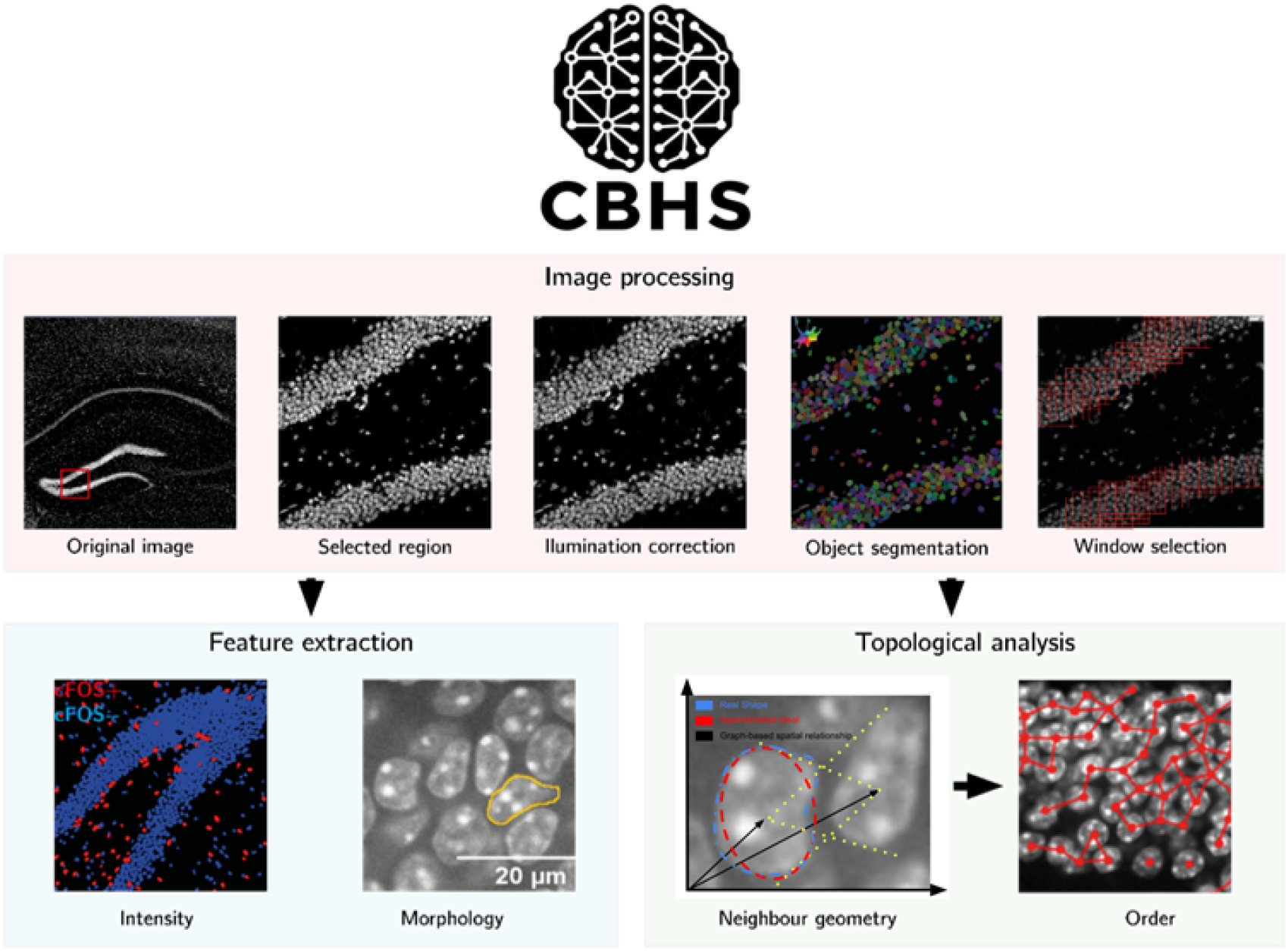
Overview of the CBHS image-analysis pipeline. Workflow from a raw confocal image through per-slice background normalisation, CellPose-based nuclear segmentation, 3D reconstruction and sliding-window selection, to feature extraction. The pipeline yields per-nucleus morphological descriptors (intensity, shape) and per-window topological descriptors (neighbour geometry, positional and orientational order).

### Intensity processing and segmentation of nuclei in high-density brain areas

To address the first issue (i), we implemented an illumination correction method that relies on erosion (*ϵ*_B_) and dilation (*δ*_B_) of the images with a structuring element B (an ellipsoid of size *q* × *q*), which enables the computation of a background correction function per slice of a *Z* -stack, independent of the heterogenously distributed cells. Afterwards, the background correction function is calculated using a Gaussian filter and finally subtracted from the original image (Table 2;Figure 2). We benchmarked the CBHS correction against cubic-spline background subtraction, which is an established approach for unevenly illuminated images (Figure 2A,B) (Carpenter et al., 2006). Then, we investigated whether this approach yields qualitative and/or quantitative improvements in terms of overall background reduction and reduced variability. A qualitative comparison on deliberately overexposed background images, together with the root mean square error (RMSE) after fitting a linear plane to the image illumination (Table 2), illustrated the difference between the two methods in background intensity and its variance across the images (Figure 2A,B). Quantitatively, the CBHS method yielded a flatter overall background intensity than both the spline correction and the uncorrected image (Kruskal–Wallis 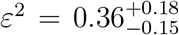, *p <* 10^−5^; CBHS median 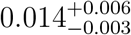 versus 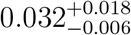 for spline and 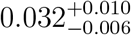 for uncorrected, *n* = 54 images ; pairwise Mann–Whitney, CBHS lower than both, *p <* 10^−5^, while spline and uncorrected did not differ, *p* = 0.83), and reduced the low-frequency spectral energy of the residual (Kruskal–Wallis 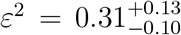, *p <* 10^−5^; CBHS median 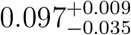 versus 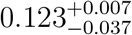 for spline and 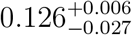 for uncorrected; CBHS lower than both, *p <* 10^−5^, while spline and uncorrected did not differ, *p* = 0.25). This metric isolates and quantifies the energy contained in low spatial frequencies of the Fourier image spectrum, components associated with gradual illumination or shading variations rather than local noise or texture. A lower low-frequency energy ratio thus indicates a more spatially uniform background illumination (Liu, et al., 2017). These data suggest that CBHS effectively corrects for variations in background intensity across regions with different cell density. Still, a non-negligible background remained after correction. Therefore, to measure intensity per-region of interest (ROI), the remaining background signals were normalized by subtracting or dividing by the equalized mean background, as summarized in Table 2. Furthermore, the background corrected signal was used for automated filtering of marker-positive cells after segmentation (Figure 2E,F,G and H).

**Figure 2:**
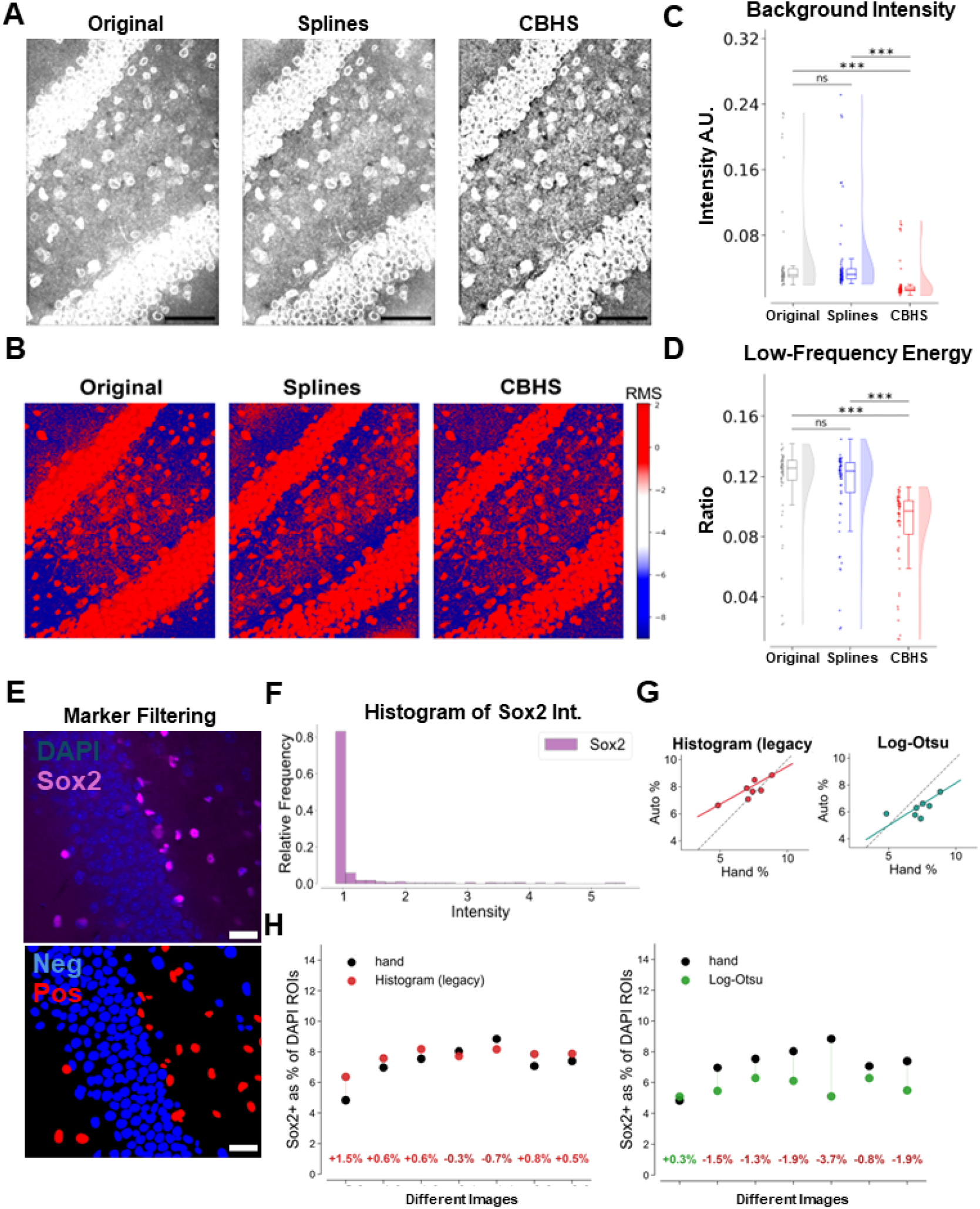
Intensity pre-processing and marker-positive cell filtering. (A) Representative overexposed Lamin B1 images illustrating background reduction after CBHS correction versus the original and spline methods (scale bar 50 µm). (B) Corresponding root-mean-square-error heatmaps of residual intensity (scale bar 50 µm). (C) Box-and-violin plots of mean background intensity, which is flattest after CBHS correction (bootstrapped Mann–Whitney; Kruskal–Wallis *ε*^2^ = 0.36; CBHS median 0.014 versus 0.032 for spline and uncorrected, *n* = 54 images; CBHS lower than both, *p <* 10^−5^). (D) The same comparison for low-frequency background energy (*ε*^2^ = 0.31; CBHS lower than both, *p <* 10^−5^). (E) DAPI/Sox2 staining in the DG with marker-positive masks (negative blue, positive red; scale bar 25 µm). (F) Representative Sox2 intensity histogram (high negative peak, long positive tail). (G) Regression of hand-annotated versus automated marker-positive percentage for the CBHS-histogram and log-Otsu algorithms (R^2^ = 0.69, RMSE 0.85%, MAE 0.60%, *n* = 7 images). (H) Per-image dot-distance plot of hand versus automated marker-positive percentage, which do not differ (paired Wilcoxon *p* = 0.19; mean difference +0.51%, *n* = 7).

Subsequently, we addressed the second challenge(ii), instance separation in densely packed tissue. Initially, we applied CellPose for segmenting the DAPI-stained nuclei, (Figure 3) (Stringer et al., 2021). Although the original CellPose Segment Anything (CPSAM) model was capable of segregating nuclei fairly well, we observed limitations in denser regions. To improve accuracy, the CPSAM model was further fine-tuned for CBHS using 4,249 hand-drawn nuclei from nine training images. The model was then benchmarked against the untuned baseline on five held-out images containing 1,446 annotated nuclei, using precision, recall, and the F-score (Figure 3A–D). Precision is the proportion of detected nuclei that matched a hand-annotated nucleus and penalizes false detections, while recall is the proportion of hand-annotated nuclei recovered by the model and penalizes misses. The F-score is their harmonic mean of two and is dominated by whichever is lower. This last property matches the cost structure of the downstream topology analysis, in which false detections (splitting a single nucleus into two) and missed nuclei propagate symmetrically into the different topological metrics used (see below). Fine-tuning consistently improved every metric: precision rose from 0.78 to 0.86, recall from 0.86 to 0.89, and F-score from 0.81 to 0.87, with the CBHS-tuned model superior on all five validation images for precision and the F-score (rank-biserial r_rb_ = 1.00) and on four of the five for recall (r_rb_ = 0.87) (Figure 3B-D) (Wilcoxon signed-rank test, *p* = 0.0625 for precision, and F-score and *p* = 0.125 for recall).

**Figure 3:**
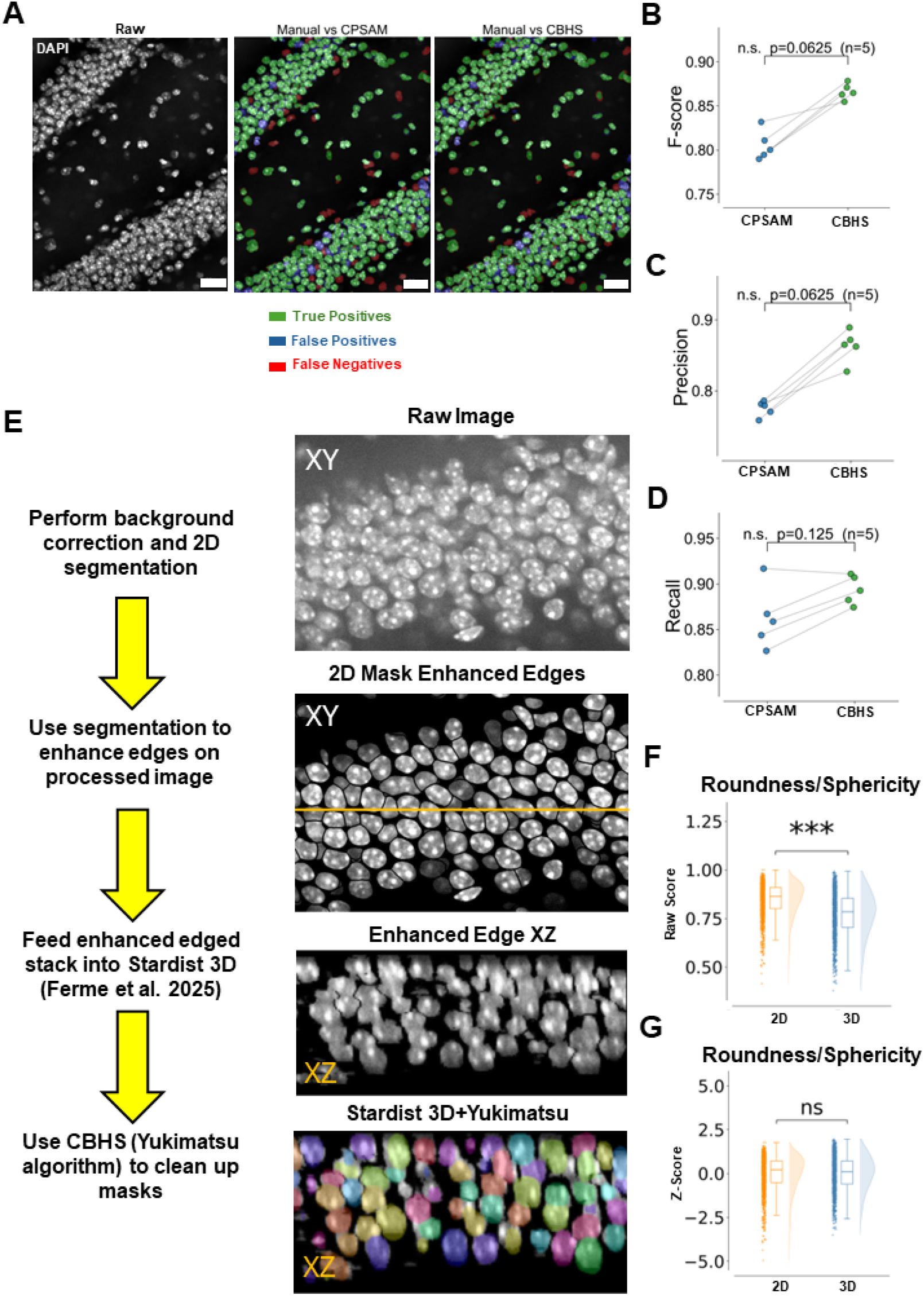
Segmentation and 3D reconstruction in high-density brain areas. (A) Raw DAPI in the DG with hand-annotated, CPSAM and CBHS-tuned masks (true positives green, false positives blue, false negatives red; scale bar 25 µm). (B–D) Per-image Wilcoxon signed-rank comparison of CPSAM versus the CBHS-tuned model across five validation images (1,446 nuclei), which improved every metric: F-score 0.81 → 0.87 (B; *p* = 0.0625), precision 0.78 → 0.86 (C; *p* = 0.0625) and recall 0.86 → 0.89 (D; *p* = 0.125). (E) Schematic and representative images of the 2D-to-3D reconstruction (mask-guided edge enhancement, Stardist 3D, Yukimatsu cleanup), shown in XY and XZ. (F) 2D roundness versus 3D sphericity on the same nuclei differ on their raw scales (box-and-violin, bootstrapped; matched-pair r_rb_ = 0.87, *n* = 1,124 nuclei), but (G) become statistically indistinguishable after z-scoring (mean bootstrap *p* = 0.44, r_rb_ = 0.09).

Following 2D segmentation, the boundaries of DAPI-positive ROIs were enhanced in each Z-stack slice based on their corresponding masks. This procedure increased the apparent separation between adjacent nuclei, thereby facilitating downstream three-dimensional instance separation. These edge-enhanced DAPI-stained nuclear image stacks were then reconstructed with Stardist 3D (Figure 3E) (Pachitariu et al., 2025; Kleinberg et al., 2022; Weigert et al., 2020). A qualitative assessment revealed that residual artifacts could distort the resulting 3D masks. To account for this, we introduced the Yukimatsu algorithm, a label-wise post-processing routine applied in two stages. The first stage restores the axially truncated nuclear caps (the issue iii): First, each labeled object is up-sampled along the *z*-axis by trilinear interpolation. After which, each object is Gaussian-smoothed, and subjected to anisotropic morphological closing with a larger structuring element along the z-axis than within the imaging plane. Lastly, they are returned to the native slice spacing by maximum-projection downsampling that preserves the recovered caps. The second stage resolves under-segmented, touching nuclei: an Euclidean distance transform seeds a watershed at the internal distance maxima, after which a set of purely geometric guardrails — a minimum child-volume (size) filter, a sphericity/elongation criterion, and a neck-thickness criterion at the shared boundary — merges spurious fragments back into single objects or filters them out, thereby suppressing false divisions.

Having obtained 3D masks (Figure 3E, Stardist 3D + Yukimatsu), we asked whether the additional computational cost of full 3D reconstruction yields more discriminative shape information than 2D analysis. We therefore compared 2D roundness with 3D sphericity (Figure 3F,G). As these two metrics are defined on different scales, their raw values differed significantly between 2D roundness and 3D sphericity on the same nuclei (after trimming the most extreme 3.45% of values per tail; bootstrapped Wilcoxon signed-rank *p <* 10^−5^, matched-pair rank-biserial 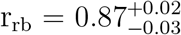; *n* = 1,124 nuclei; Figure 3F). However, after standardization to *z*-scores, the distributions of the two factors were statistically indistinguishable under the size-matched bootstrap scheme (Figure 3G, the mean bootstrapped *p* = 0.44, the matched-pair effect size 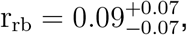, *n* = 1,124; see also Materials and Methods for Statistics for the usage of bootstrapped scheme). Thus, 2D roundness preserves the same relative shape information as 3D sphericity. Based on this observation, all subsequent analyses were performed on 2D slice images.

Once the background correction and segmentation had been validated, we tested whether cell type-specific marker-positive cells can be identified automatically. In the case of nuclear markers, the mean intensity within each ROI (nuclei) divided by the corrected background intensity is calculated (Table 2). In the case of cytoplasmic markers, the mean intensity of the pixel shell immediately outside each ROI divided by the background intensity is calculated (Table 2). In the hippocampus, many of the markers used, for instance Sox2, a transcription factor for adult neural stem/progenitor cell marker, shows a stereotypical intensity distribution across different cells (Figure 2E,F): a sharp first peak (the marker negative cells) followed by a long right tail, reflecting heterogeneous expression levels within an otherwise uniformly positive population. Several thresholding strategies have been described for such distributions (Jumiawi and El-Zaart., 2022). Here, we evaluated log-Otsu thresholding alongside an in-house developed algorithm, called “histogram”. The histogram locates the first high peak and shifts a pre-specified number of *z*-score standard deviations in the positive direction to set the cut-off; the step size can be calibrated once for a given marker class and then held constant across conditions. This algorithm is specifically designed for already background-corrected ROIs, as the *z*-score based distribution of values follows the same pattern, allowing for consistent cross-comparisons of different cell-types, whose intensity was corrected with the same method.

To validate both thresholding methods, the automated Sox2-positive percentage was compared with hand-counted percentages across seven images drawn from seven distinct DGs. Agreement metrics indicated moderate predictive accuracy (Figure 2G, R^2^ = 0.69, Root Mean Squared Error (RMSE) = 0.85%, and Mean Absolute Error (MAE) = 0.60% (*n* = 7 images)). However, the residual error (RMSE & MAE) motivated a direct paired comparison between hand and automated counts across the seven images, which revealed no statistically significant difference between hand and automated counts (Figure 2H, paired Wilcoxon signed-rank *p* = 0.19; paired t, *p* = 0.12; mean difference 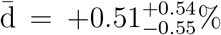; *n* = 7). Together, the background correction, segmentation of nuclei and cell-type specific marker filtering, lay the foundation for the following investigation of topological order and nuclear shape across the different hippocampal subfields.

### Positional and orientational order across hippocampal subfields

Topological and geometric cellular organization can be assessed at the mesoscale through a range of metrics, all of which ultimately describe the distances between nuclei, their angular arrangement relative to one another, and the shape of the individual nuclei. To estimate the cellular packing order from each brain region, sliding square windows were applied that partition the field into equally sized rectangles. In this way, the structural arrangement and orders can be compared in the same dimension (Figure 1). Within this framework, in addition to basic histological features, we examined the positional and orientational order of nuclei in the brain. Positional order informs about the spacing and chance of occurrence of other cells as a function of separation distance, while orientational order informs about the shape alignment towards a bond reference-orientation (Figure 4A). Both were computed in 2D across several slices of the same *Z* -stack. Positional order was quantified using the radial distribution function *g*(*r*) together with the associated two-point entropy (*S*_2_), whereas orientational order *S* was derived from the local nematic order parameter (Table 2; Figure 4A)(Kim et al., 2024; Ferme et al., 2025).

**Figure 4:**
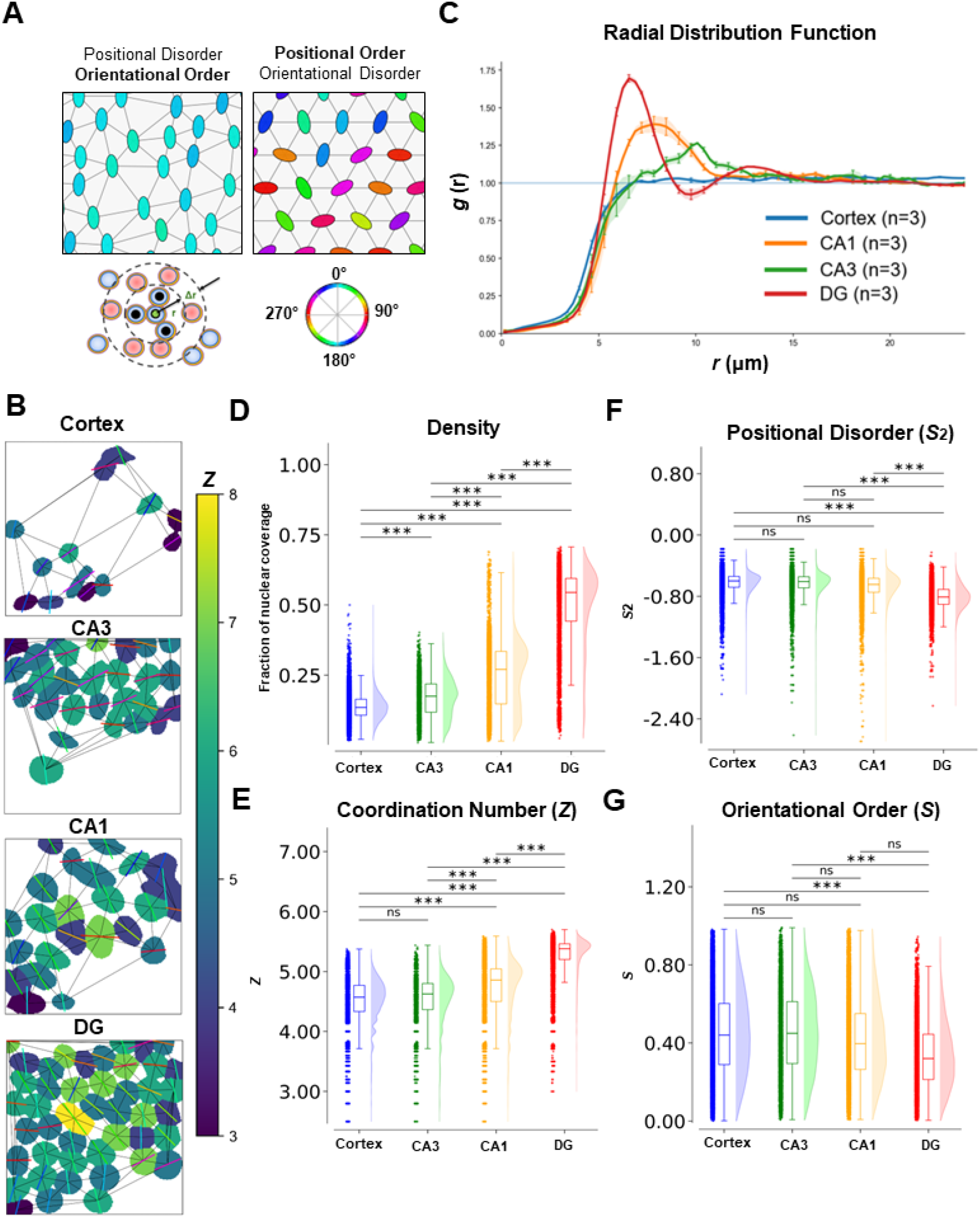
Positional and orientational order across hippocampal subfields. (A) Schematic of positional order (from the radial distribution function) and orientational order (from the local nematic order parameter). (B) Representative masks and coordination-number heatmaps for cortex, CA1, CA3 and DG. (C) Mean radial distribution function *g*(*r*) per subfield (line width = SEM across all windows from three mice), with the highest and earliest first peak in the DG. (D– H) Per-window box-and-violin plots with bootstrapped pairwise Mann–Whitney for nuclear density (D; Kruskal–Wallis *ε*^2^ = 0.40; DG densest at coverage 0.55 versus cortex 0.13; DG-versus-cortex Cliff’s *δ* = 0.93), coordination number *Z* (E; *ε*^2^ = 0.35; DG 5.38; *δ* = −0.87), positional order *S*_2_ (F; *ε*^2^ = −0.19; DG 0.81, the lowest *S*_2_ and thus strongest positional order; *δ* = − 0.69) and orientational order *S* (H; *ε*^2^ = 0.04; DG 0.32, least aligned; *δ* = −0.32). Window counts range from *n* = 9,000 (DG) to 25,925 (cortex).

Across the DG, CA1, CA3 and cortex, we first compared simple histological features, including nuclear density and inter-nuclear neighbour centroid-centroid distance(INCD). Distinct density of nuclei and INCD emerged across regions. The DG exhibited the smallest INCD and the highest density of nuclei, as defined by the nuclear coverage over the window size. (Table 2; Kruskal–Wallis across subfields, nuclear density 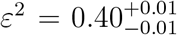, *p <* 10^−5^ ; INCD *p <* 10^−5^; DG densest, at coverage 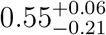 *(n* = 4,275 windows) and shortest spacing 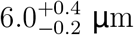; CA1 next (coverage 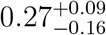, *n* = 7,650; spacing 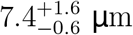), then CA3 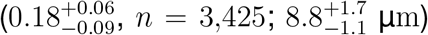 and the cortex 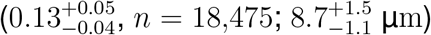 ). The greatest differences were observed in the density of nuclei and the INCD between the DG and the cortex (Figure 4C,D, Cliff’s 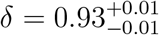 for density, 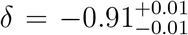 for distance, pairwise Mann–Whitney tests *p <* 10^−5^,). CA1 showed the second-smallest distances and the second-highest density, followed in turn by CA3 and the cortex (Figure 4C,D). Motivated by the quantitative differences in density and distance, we then analysed the spatial relationships of neighbouring nuclei using *g*(*r*). Consistent with this density, the DG also displayed the strongest positional order, reflected in the highest first peak of the *g*(*r*) and in its being closest to the reference nucleus. The strong positional order is also well represented by the per-subfield mean *g*(*r*) curves. The strongest positional order of DG is followed by CA1, and CA3 (DG first peak *g*(*r*) ≈ 1.7 at *r* ≈ 6.5 µm; CA1, ≈ 1.4 at ≈ 7 µm; CA3, ≈ 1.3 at ≈ 8 µm). The *g*(*r*) of the cortex was essentially flat, indicating near-random, homogeneous spatial profile (*g*(*r*) ≈ 1.05; Figure 4C). Based on the *g*(*r*), which represents the order of cellular alignment in the tissue, we also calculated *S*_2_, the integral pair-correlation measure that estimates pairwise entropy in the given structure. We found that DG exhibited the lowest *S*_2_ followed by CA1, CA3 and the cortex (Figure 4F). These data suggest that the DG inherently possesses the highest level of positional order (mean DG, 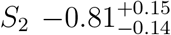, *n* = 9,000 windows; CA1, 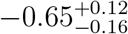, *n* = 11,600; CA3, 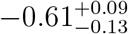, *n* = 5,150); and Cortex, 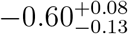, *n* = 25,925); Kruskal–Wallis 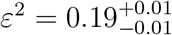, *p <* 10^−5^, with DG versus each subfield significant at the bootstrap threshold (Figure 4F; bootstrapped Mann–Whitney, proportion of resamples significant = 1.00 in every case; DG versus cortex median *p <* 10^−5^, Cliff’s 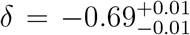; DG versus CA3 median *p <* 10^−5^; DG versus CA1 median *p <* 10^−5^, cortex versus CA3 *p* = 0.45, CA3 versus CA1 *p* = 0.067, cortex versus CA1 *p* = 0.016, all n.s. at the bootstrap threshold). As only the DG shows a significant difference in positional order, a more detailed investigation of the exact breakpoint of order over density is warranted.

To further characterize the surrounding spatial relationships between nuclei within each brain region, we measured the coordination number *Z*, which is defined as the mean number of neighbours per nucleus (Figure 4B,E). Consistent with the higher density of nuclei across the window and shorter inter-nuclear spacing, the DG exhibited the highest *Z* value, followed by CA1, CA3, and the cortex (Kruskal–Wallis 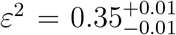, *p <* 10^−5^; DG, 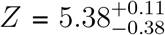, *n* = 9,000 windows; CA1. 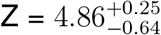, *n* = 11,600) ; CA3 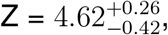, *n* = 5,150); Cortex 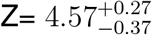, *n* = 25,925). The difference between the DG and the cortex is the greatest (Cliff’s 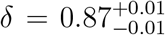). The coordination numbers of the DG and CA1 differ from every other subfield, but not from those of the cortex and CA3 (Figure 4E, (*p* = 0.34). These data suggest that, in the DG, each nucleus is not only closer to others, but also that each nucleus has a higher number of potentially interacting neighbours. Following the difference in spatial cellular alignment among brain regions, we next measured orientational order, which represents the cellular alignment towards a shared reference orientation. Compared to positional order, orientational order exhibited an opposite trend, with the least aligned nuclei in the DG and the next least aligned in CA1, while CA3 and the cortex were the most aligned and statistically indistinguishable (Kruskal–Wallis 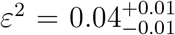; DG, *p <* 10 ^−5^ ; 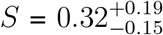,*n* = 9,000 windows; CA1 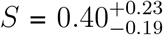, *n* = 11,600; cortex 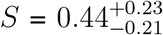, *n* = 25,925; CA3, 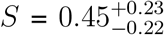, *n* = 5,150). The effect size between the DG and the cortex is moderate (Cliff’s 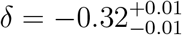), and only the DG versus cortex (*p* = 1.2 × 10^−4^) and DG the versus CA3 (*p <* 10^−5^) exhibited significant differences (Figure 4G). Additionally, the effect size for orientational order was considerably smaller than that observed for positional order (rank effect size *ε*^2^ = 0.19 for positional (*S*_2_) versus *ε*^2^ = 0.04 for orientational (*S*) order, a roughly fourfold smaller effect, mirrored by the DG-versus-cortex Cliff’s 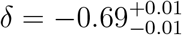 for *S*_2_ versus 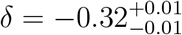 for *S*; *n* = 51,675 windows). These data suggest that orientational order, compared to positional order, might not carry the same wealth of information. The local nematic order relies on a shared reference orientation. Unfortunately, establishing such a reference orientation is noisy for more circular nuclei.

### The relationship between order and density across hippocampal subfields

Having established that order and density co-vary across subfields quantitatively, we next examined how order scales with density within each region by applying a series of fits to the full population of sliding windows (Figure 5A–C). We first used linear fits of positional order against density and found that only the DG exhibited a clear monotonic relationship. The higher the density, the lower the *S*_2_ levels, indicating an increase in positional order at higher nuclear coverage (Figure 5A,B). On the other hand, the relationship between *S*_2_/*S* and cell density in CA1, CA3, and the cortex, were essentially flat (Figure 5A; Ordinary Least Squares (OLS) of *S*_2_ on density: DG 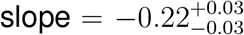, R^2^ = 0.05, *p <* 10^−5^, *n* = 4,275 windows; CA1 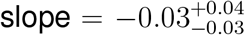, *p* = 0.10, *n* = 7,650; CA3 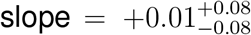, *p* = 0.83, *n* = 3,425, both n.s.; cortex slope 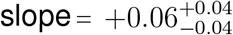, *n* = 18,475). This data suggests that an increase of positional order with density only happens in the DG, which is consistent with the findings of the previous subsection. By contrast, for orientational order, all subfields showed a negative association between *S* and density (Figure 5A, OLS of *S* on density, all slopes negative and significant: DG slope = − 0.30,R^2^ = 0.08, *n* = 4,275; CA1 slope = −0.39, R^2^ = 0.06, *n* = 7,650; CA3 slope = −0.27, R^2^ = 0.01, *n* = 3,425; cortex slope = −0.44, R^2^ = 0.01, *n* = 18,475, all *p <* 10^−5^). These data replicate the findings from Figure 4, illustrating how the relationship between density and positional order is strongest in the DG, while the orientational order carries little weight to our understanding of how order emerges from density.

**Figure 5:**
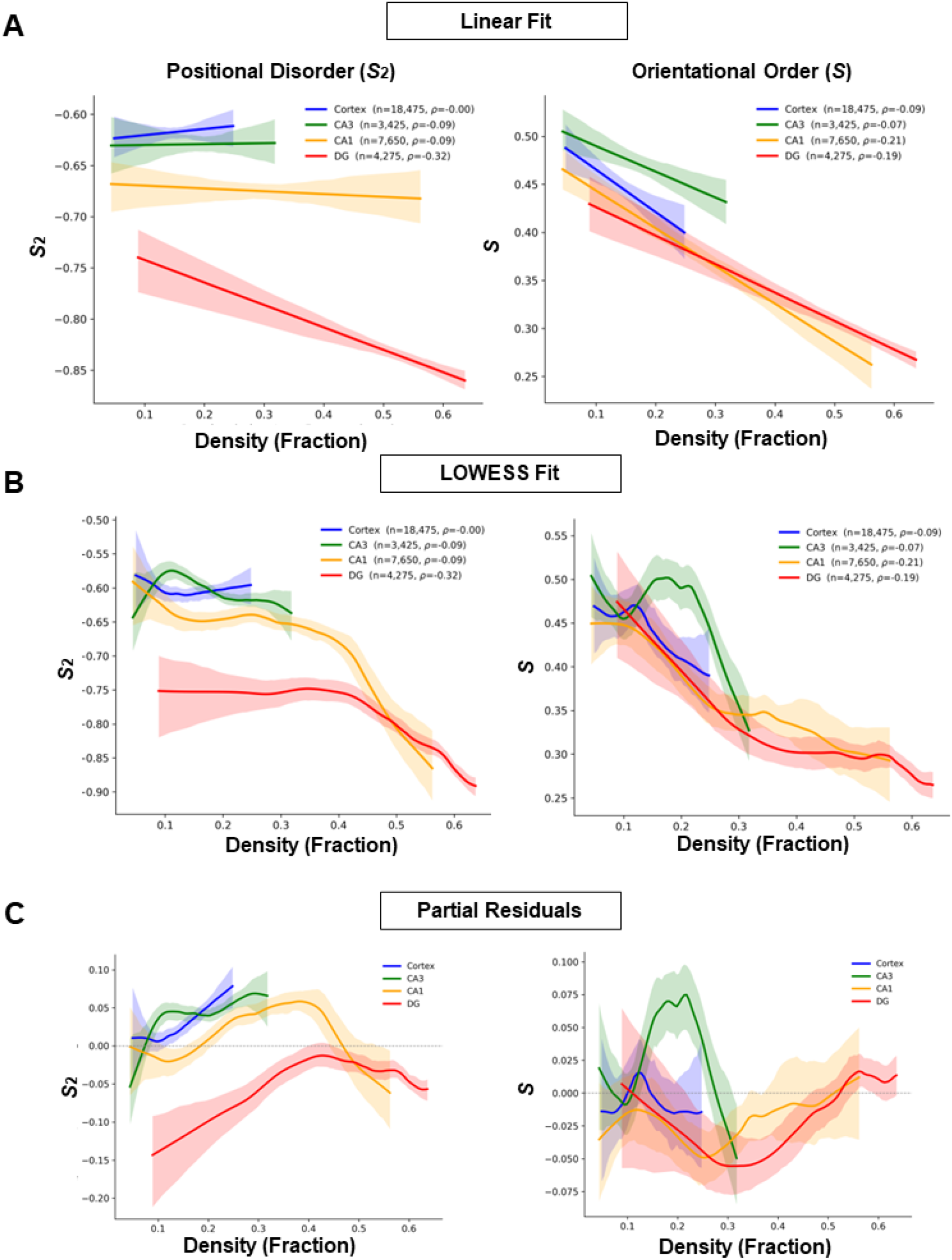
Relationship between order and density. (A) Linear fits of positional order (*S*_2_) and orientational order (*S*) against nuclear density across all windows from cortex, CA1, CA3 and DG (band = 95% CI); ordinary-least-squares regression of *S*_2_ on density is significant only in the DG (slope −0.22, R^2^ = 0.05, *p <* 10^−5^). (B) LOWESS fits reveal a segmented breakpoint in the *S*_2_–density relationship at a density of 0.39 (DG) and 0.35 (CA1), beyond which positional order rises steeply (top-quartile CA1 slope −0.89). (C) LOWESS fits on partial residuals corroborate this breakpoint structure; orientational order instead decreases with density in every subfield (Spearman, DG *ρ* = −0.19). In all panels the band width denotes the 95% confidence interval of each fit, with *n* = 3,425–18,475 windows per subfield.

Next, we investigated whether there is a clear breakpoint at which order emerges from increasing density, where cells become ordered. Because these linear fits assume a single global slope, we therefore applied locally weighted (LOWESS) fits to capture any density-dependent changes in the relationship between cellular order and density (Figure 5B). Indeed, the LOWESS curves indicated a non-linear relationship between density and positional order. Positional order increased above a breakpoint of approximately 0.4 nuclear coverage in the dense DG and CA1 subregions. Spearman correlation of *S*_2_ on density over the full range indicated that only the DG shows a robust association between the positional order and density (*ρ* = −0.32, *p <* 10^−5^, *n* = 4,275). In CA1 and CA3, their association was mild (CA1 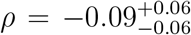, *n* = 7,650; CA3 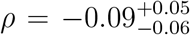, *n* = 3,425; neither significant under subsample bootstrapping) and no association was apparent in the cortex (*ρ ≈* 0, *p* = 0.94, *n* = 18,475). A segmented regression identified a significant breakpoint in the *S*_2_–density relationship in every subfield (segmented vs. linear *p <* 10^−5^), positioned at a nuclear density of 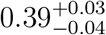 in the DG and 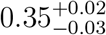 in CA1. However, above the DG and CA1 breakpoints the slope reversed sign (Figure 5B; DG +0.27 → − 0.57; CA1 +0.15 → −1.00). Beyond the break point, higher-density windows in the DG/CA1 exhibited decreasing *S*_2_, indicating greater positional order with increasing density. To confirm this observation using a linear regression fit, the OLS fit was applied to the top quartile of CA1 density windows, indicating a steep and robust negative correlation between density and positional order within that range (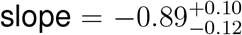, R^2^ = 0.18, *p <* 10^−5^, *n* = 1,913 windows). In contrast, orientational order decreased steadily with density across all subfields and, interestingly, plateaued at high densities near the same nuclear density of ≈ 0.4 in the DG and CA1 (Figure 5B, Spearman correlation of *S* on density: CA1, *ρ* = −0.21, *p <* 10^−5^, *n* = 7,650; DG, 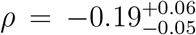, *n* = 4,275; cortex, 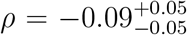, *n* = 18,475; CA3, 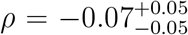, *n* = 3,425; the latter two are not significant under subsample bootstrapping). The orientational decline likewise showed a significant breakpoint (DG at 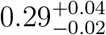, slope −0.81 → −0.15; CA1 at 0.27; both *p <* 10^−5^), with the residual decline flattening above ∼0.3; Figure 5B). The CA3 curve was notable in rising slightly before declining in S (descriptive, LOWESS curve shape; CA3 overall Spearman 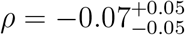, a weak association; Figure 5B). Together, these results suggest that spatial order arises at a certain density in a cross-subfield coherent manner.

Finally, to assess whether the linear term is adequately specified, we utilized partial-residual plots, which isolate the density contribution after accounting for the remaining model terms. The partial residuals showed that positional order increased with density up to approximately 0.4 before decreasing again (partial residuals; Spearman of residual versus density robustly positive in CA1 (*ρ* = 0.21, *p <* 10^−5^, *n* = 7,650), weakly positive in the cortex (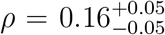, *n* = 18,475) and CA3 (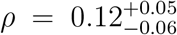, *n* = 3,425), and flat in DG (*ρ* = 0.02, *p* = 0.13, *n* = 4,275). The turning point near 0.4 can be seen from the curve in Figure 5C. These results suggest that the linear term is adequately specified, but only before and after the breakpoints. In most regions, orientational order displayed the opposite pattern, decreasing to a minimum near a coverage of 0.3 before rising again. CA3 was again the exception, showing a sharp rise followed by a fall that paralleled the positional-order trend (see Figure 5C for orientational partial residuals; CA3 shows a positive residual offset and CA1 a negative one, with near-zero offsets in the cortex and DG). Consistent with this reversal relationship, the Spearman of the orientational residual versus density was robustly positive in DG (*ρ* = 0.12, *p <* 10^−5^, *n* = 4,275) — the one region that had been flat for positional order — weakly positive in CA3 (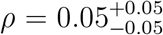, *p* = 0.005, *n* = 3,425), and flat in cortex (*ρ* = − 0.01, *p* = 0.07, *n* = 18,475) and CA1 (*ρ* = 0.01, *p* = 0.24, *n* = 7,650), the region that had carried the strongest positional trend. Intriguingly, only subfields that reach the density breakpoint show an increase in order. Nevertheless, once reaching the breakpoint, both CA1 and DG show a similar slope of increased positional order. Intriguingly, orientational order does not have the same breakpoint. This is likely because nuclear shape dependent orientational differences arise in a subfield- and cell type-specific, rather than in density-associated manner. This observation raises the question of whether the shape of the nuclei themselves could become altered under the physical constraints of their neighbours.

### The relationship between density and nuclear shape

Our mesoscopic investigation of nuclear organization in the brain raised the possibility of nuclear deformation depending on the given nuclear density (Figure 5). In fact, nuclei in high-density regions in the DG alter their shape depending on neighbouring cells (Figure 6A). This relationship may contribute to the coupling between order and density described above. Consistent with this idea, at the window level, the average nuclear aspect ratio increased with density across all subfields (Figure 6B; Spearman, CA1 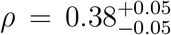, *n* = 7,650; CA3 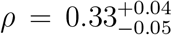, *n* = 3,425 (both *p <* 10^−5^); DG 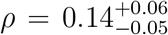, *n* = 4,275; cortex 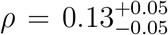, *n* = 18,475), indicating that nuclei became more circular (with aspect ratio approaching 1) in more crowded (higher density) regions. In the DG, however, this increase levelled off, with the average aspect ratio reaching a plateau above a density of approximately 0.45 (Figure 6B). This suggests that at high densities, nuclei might undergo different shape constraints after a given breakpoint.

**Figure 6:**
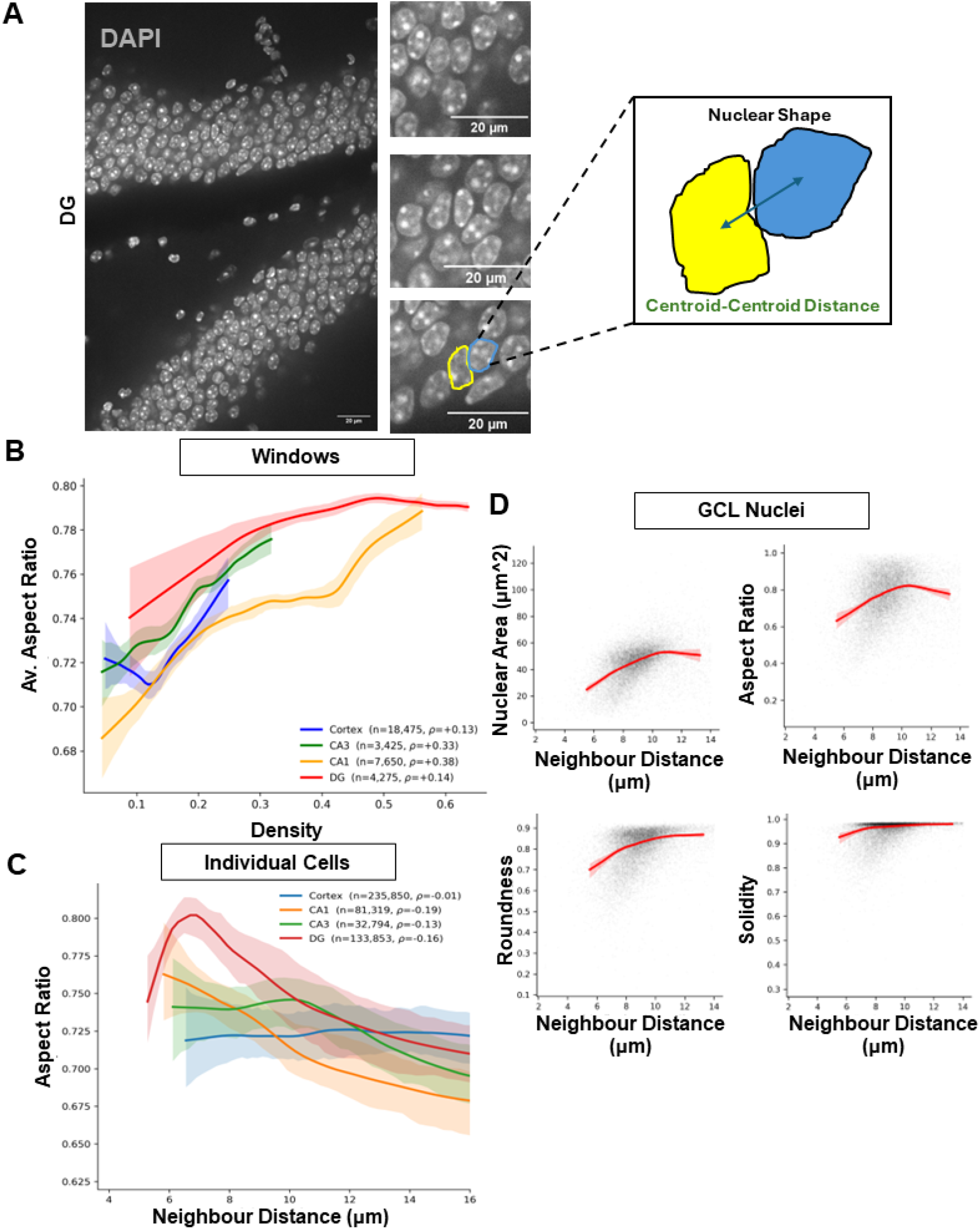
Density and nuclear shape in the dentate gyrus. (A) Raw DAPI from the DG with cropped regions highlighting nuclear deformation between granule cells, and a schematic of nuclear shape and centroid-to-centroid distance (scale bar 20 µm). (B) Window-averaged aspect ratio increases with density in every subfield (LOWESS, 95% CI; Spearman CA1 *ρ* = 0.38, CA3 0.33, DG 0.14, cortex 0.13). (C) Single-nucleus aspect ratio decreases with the mean centroid-to-centroid distance to the three nearest neighbours (LOWESS, 95% CI; CA1 *ρ* = −0.19, DG −0.16, CA3 −0.13; cortex ≈0). (D) Area, aspect ratio, roundness and solidity of GCL nuclei all rise with neighbour distance before plateauing (LOWESS fit with per-cell dots; Spearman area *ρ* = 0.47, roundness 0.42, solidity 0.42, aspect ratio 0.30; *n* = 15,743 nuclei).

To resolve this issue at the single-nucleus level, we examined the relationship between the aspect ratio of individual nuclei and the mean centroid-to-centroid distance to their three nearest neighbours (Figure 6C). The association between shorter neighbour distances and higher aspect ratios was strongest in the CA1 and DG, and weakly in the CA3 (Spearman over individual nuclei: CA1 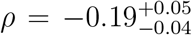, *n* = 81,319; DG 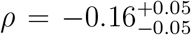, *n* = 133,853; CA3 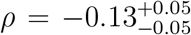, *n* = 32,794)). These associations are small and their CIs exclude zero but they fall below the bootstrap-significance threshold despite the very large *n* (Figure 6C), whereas the cortex showed essentially no relationship (*ρ* = −0.01; *n* = 235,850 nuclei). In the DG, this trend reversed at the lowest distances, with the aspect ratio decreasing again below approximately 6.5 µm (Figure 6C). These data are consistent with our idea that at very high cellular density there might be additional shape altering constraints.

To characterize how nuclear shape is altered in high-density regions, we quantified the area, aspect ratio, roundness, and solidity of granular cell layer (GCL) nuclei as a function of neighbour distance (Figure 6D). All four parameters exhibited lower values at shorter neighbour distances rising before plateauing at approximately 9 µm of neighbour distance (Figure 6D, Spearman across *n* = 15,743 GCL nuclei, all positive versus neighbour distance: area 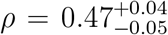, roundness 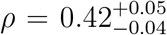, solidity 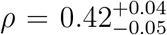, aspect ratio 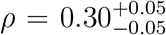; all fits *p <* 10 ^−5^ ). The aspect ratio was the only descriptor not to reach a plateau, decreasing slightly again at larger distances (Figure 6D, quadratic term fit *p <* 10^−5^; slope = −0.010, *p <* 10^−5^). Together, these data illustrate how at high-density nuclear shape reaches a breaking point, at which nuclei become less circular again, after facing additional constraint from their surrounding environment. Importantly, the GCL consists of several cell types, the majority of which are dentate granule cells. Different neural cell-types are known to have different deformability (Lu et al., 2006). Hence, the relationship between the local environment and nuclear shape exists in a cell type-dependent manner.

### Nuclear shape across neural cell types

Our quantitative analyses revealed that density of nuclei and order are interconnected in a brain region-specific manner. While we analyze all DAPI-positive nuclei together, different neural cell types are known to differ in nuclear shape and deformability (Lu et al., 2006). This raised the question of whether the nuclear shape of distinct cell types is differentially related to the local environment (Figure 7A). We therefore compared Ctip2- and/or NeuN-positive neurons with GFAP- and/or S100β-positive astrocytes across low- and high density windows in the hippocampus, including the DG, CA1 and CA3. Each nucleus was assigned to a high- or low-density local environment according to the centroid-to-centroid distance to its nearest neighbour: nuclei whose nearest neighbour lay within 12 µm were classified as high-density, whereas those with a nearest-neighbour distance of 12 µm or greater were classified as low-density. After nuclear segmentation and cell type annotation, nuclear morphological information and local environmental information are extracted using CBHS. To compare the influence of cellular density, a principal component analysis (PCA) of all of the nuclear shape factors listed in Table 2, is computed separately for low- and high-density local environments. In both density regimes, neurons and astrocytes occupied a single, heavily overlapping region of shape space rather than separable clusters, with the first two principal components accounting for 72.6% of the shape variance at low density and 71.8% at high density (Figure 7B,C, *n* = 27,936 low- and *n* = 107,664 high-density nuclei). Cell type was a statistically significant but minor source of shape variance (PERMANOVA *p <* 0.001, both densities), explaining only under 5% of the multivariate variance (*R*^2^ = 0.03–0.05 across both densities, the PC1–PC2, and the full 15-factor spaces), with the neuron and astrocyte centroids-space separated by only 0.40–0.52×, which is the usual range of variation within the type. However, the cell types differed more in their shape variability than in their mean shape. Astrocytes occupied a larger region of shape space than neurons (PERMDISP *p <* 0.001 at both densities), with an astrocyte dispersion 1.10–1.13× that of neurons across the full multivariate space, and 1.2–1.3× on the one-dimensional shape-irregularity composite. This dispersion difference is established by the balanced-subsample PERMDISP above and corroborated by Levene’s test, *W* = 453 at low and *W* = 818 at high density, whose extreme nominal *p*-values reflect the very large *n* (see Figure 7B,C). The two neuronal markers, Ctip2 and NeuN, did not differ in shape variability at low density (PERMDISP *p* = 0.85). These data suggest that the majority of cells share a similar nuclear shape across cell types. The difference arises from the extremes that each cell type reaches. Hence, nuclear shape is not stereotypical according to the present analysis.

**Figure 7:**
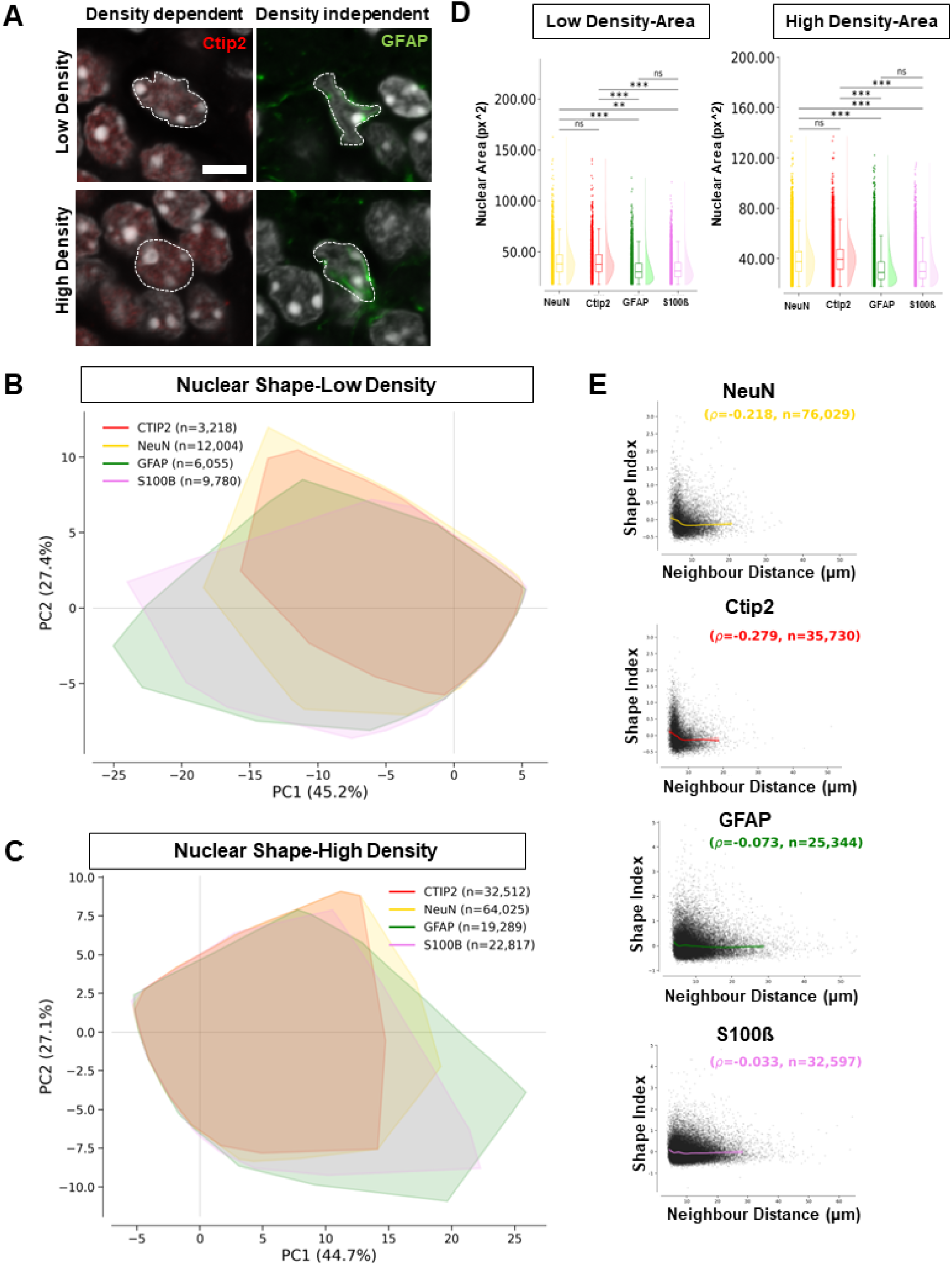
Nuclear shape across neural cell types. (A) Low- and high-density DG crops for a CTIP2-positive neuron and a GFAP-positive astrocyte (scale bar 5 µm). (B,C) Principal-component analysis (convex hulls) of nuclear shape for CTIP2-, NeuN-, GFAP- and S100β-positive nuclei at high (B) and low (C) neighbour distance; the first two components explain 71.8% (high) and 72.6% (low) of the shape variance, and cell type is only a minor source of variance (PERMANOVA *R*^2^ = 0.03–0.05), although astrocytes are more dispersed than neurons (1.1–1.3 ×, PERMDISP). (D) Nuclear area per cell type and local-density environment (box-and-violin, bootstrapped pairwise Mann–Whitney): neuron nuclei are larger than astrocyte nuclei (37.5 versus 30.9 px^2^ at low density; Cliff’s *δ* = 0.26–0.30). (E) The composite shape index varies with neighbour distance only for neurons (LOWESS fit with per-cell dots; Ctip2 *ρ* = −0.28, NeuN −0.22) and not astrocytes (GFAP −0.07, S100β −0.03); *n* = 27,936 (low) and 107,664 (high) nuclei.

We next compared the nuclear areas between the two cell-types. Neurons had larger nuclei than astrocytes, consistent with previous reports (Garcia-Cabezas et al., 2016; Das Gupta et al., 2023). This held true at both low and high density (bootstrap Kruskal–Wallis *p* ≈3 × 10^−4^ at low and *p <* 10^−5^ at high density), with a small-to-moderate effect in each case (Cliff’s 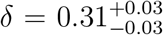 at low density; *δ* = 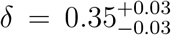 at high density). The nuclear area of neurons was 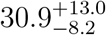 (*n* = 12,101) versus 30.9^+13.0^px^2^ for astrocytes (*n* = 15,835) at low density, and 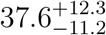 for neurons (*n* = 65,558) versus 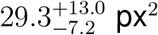 for astrocytes (*n* = 42,106) at high density. In pairwise Mann–Whitney tests, the nuclear areas of neurons were significantly larger than both astrocyte markers. However, differences within the same cell types (NeuN–Ctip2 and GFAP–S100β) were not significant (Figure 7D).

Finally, to investigate, whether the dependence of nuclear shape on the local density is cell-type specific, we defined a composite shape index from the individual shape metrics (Table 2) and compared its LOWESS fit with that of neighbour distance. By comparing the shape index and the distance from the neighbour, we found that only the nuclear shapes in the neurons showed a dependence on the local environment, with the shape index increasing at shorter distances from the neighbour (Figure 7E, Ctip2 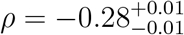, *n* = 35,730), NeuN 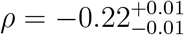, *n* = 76,029, both *p <* 10^−5^ ;). Conversely, astrocytes showed no such relationship (Figure 7E; GFAP 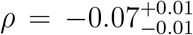, *n* = 25,344; S100β 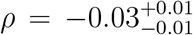,*n* = 32,597, the latter not significant by Pearson). These observations suggest two nonexclusive possibilities. The first possibility is that the relationship between nuclear shape and density is indeed cell-type specific. The second is that this relationship could arise from the large inherent variability in astrocytes, whereby the constraints of the local environment are insufficient and might not be enough to yield additional deformation.

### Equations

All equations are displayed and referenced in Table 2.

## DISCUSSION

In the present study, to investigate the mesoscale structure of the hippocampus, we developed an automated image-analysis pipeline, termed CBHS. Its background-correction step reduces background variability across the image and thereby enables reliable per-ROI background normalization. CellPose-based refined nuclear segmentation achieves automated delineation comparable to hand-drawn ROIs, while Stardist provides high-quality 3D reconstruction. However, our analyses revealed that 3D reconstruction adds little information beyond 2D for group-based comparisons of nuclear shape. After segmenting each nuclei, identifying marker-positive cells is essential for understanding how individual cell types contribute to the mesoscale structure. The histogram-based filtering implemented in CBHS achieves an accuracy comparable to that of manually-annotated ROIs. Together, these components allow CBHS to perform automated analysis of high-quality images from densely packed brain regions, such as the hippocampus.

Building on this pipeline, we quantitatively characterized the mesoscale organization of the hippocampus through its positional and orientational order. Our findings revealed that the cellular arrangement of the hippocampal subfields differ intrinsically, with the denser regions, most notably the DG, being the most positionally ordered. Intriguingly, the orientational order followed the opposite trend, although with a much lower effect size. This suggests that local nematic order is not a reliable indicator of the nuclear organization in the hippocampus, probably because its orientation is also dependent on nuclear shape, which ultimately reduces effect sizes to negligible levels. Notably, the increase in positional order emerged above a statistically resolved density breakpoint, located near ≈0.4 in the denser DG and CA1 area. This led us to the hypothesis, that above this breakpoint, the density-driven constraint of cells on each other drives order. In line with this idea, examining nuclear shape in relation to the local environment revealed a scale-dependent picture: the window-averaged nuclear shape became more circular at higher density, yet individual nuclei were smaller, less rounded, and more elongated the closer their neighbours were. This apparent paradox can be reconciled by recognizing that the two measurements are aggregated at different scales and conditioned on different quantities. The window average is dominated by the majority of moderately spaced nuclei, which pack more uniformly, and therefore appear rounder on average, as a region fills. In contrast, the elongation is concentrated in the minority of nuclei in closest contact, where direct crowding is greatest. This dependence on the local environment prompted us to ask whether the relationship between environment and nuclear shape is cell type specific. By comparing neurons with astrocytes, which themselves differ fundamentally in the variance of their nuclear shape, we found that only neurons exhibited a dependence of nuclear shape on the local environment (Figure 7). Several features of these findings strike us as particularly intriguing. First, the emergence of the cellular packing order above a statistically resolved density breakpoint, ≈ 0.4 in the more ordered DG and CA1 regions, is reminiscent of the density- or packing-driven transitions described for jammed and active matter, in which global spatial order only arises once a critical local density is exceeded (Kim et al., 2024; Zaccone, 2025). This breakpoint (a nuclear coverage of ≈ 0.4 in these regions) sits well below the saturating coverage of the tissue (≈ 0.6) suggests that hippocampal nuclear arrangements are not merely passively packed. They may reorganize their mesoscale structure once crowding becomes sufficient to constrain neighbouring nuclei, a behaviour that would not be expected from cells positioned independently of one another. Second, the cell-type specificity of the shape–environment relationship is equally striking: although astrocytes display greater intrinsic variance in nuclear shape, only neurons adapt their nuclear shape with the local environment. This dissociation argues against the coupling being merely a trivial consequence of denser regions containing systematically different cells, and instead points to a genuine, cell type-restricted responsiveness of neuronal nuclei to their surroundings. Taken together, these observations support the idea of a mechanical interaction between neighbouring nuclei that contribute to the emergence of mesoscale order. However, we emphasize that this is only one of several possible explanations. The same correlations could arise from cell-intrinsic or developmental programmes that co-vary with regional density, or from differences in baseline deformability between neurons and astrocytes, even in the absence of any direct mechanical coupling. Distinguishing among these possibilities will require perturbative or longitudinal approaches rather than the static, observational data presented here. Nevertheless, because nuclear shape and cellular deformability have been implicated in neuronal function and gene regulation (Wittmann et al., 2009), the hypothesis of a mechanical contribution to mesoscale order remains an attractive and experimentally feasible hypothesis for future investigations. The phenotype descriptions presented in this study could now be used to investigate mesoscale differences across disease conditions, as well as for functional investigations, for example in an evolutionary context.

## Limitations of the study

CBHS is intended as an image-analysis tool for investigating the mesoscale structure of the hippocampal subfields. Although it enables accurate quantification of topological order, marker intensity, and nuclear shape, the present study establishes neither a causal relationship between these features nor their biological relevance. It likewise does not address whether mesoscale order is able to adapt across conditions such as ageing or disease; the organisation reported here may instead represent a static, phenomenological description of these neuronal ensembles. Resolving these questions, and in particular distinguishing functional order from incidental structure, will require future studies.

## Lead contact

Requests for further information and resources should be directed to and will be fulfilled by the lead contact, Prof.Dr.Tomohisa Toda.

## Data and code availability

- Images can be made available upon request to the leading author
- All original code has been deposited at Gitlab under the permalink https://gitlab.mpcdf.mpg.de/konmi/image-analysis-ui/-/tree/e1eea9740df5e9c3e5e0a185510e0f82145f9623 and is publicly available as of the date of publication.
- Any additional information required to reanalyze the data reported in this paper is available from the lead contact upon request.

## ACKNOWLEDGMENTS

This work was funded by Deutsche Forschungsgemeischaft[470322152 - TO1347/3-1 (T.T.); 497658532 - TO1347/4-1 (T.T.); 507965872 - TO1347/5-1 (T.T), 563413271 -TO1347/6-1 (T.T.), CRC1540 Exploring Brain Mechanics - 460333672 (T.T., S.M., J.G.)], Schram Foundation (T.T.), the European Research Council (ERC-2018-STG, 804468 EAGER; ERC-2023-COG, 101125034 NEUTIME (T.T.)), the Interdisciplinary Center for Clinical Research (IZKF) Erlangen (P162. T.T.), the Emmy Noether Programme of the German Research Foundation (project 455449456 (J.K.)) and the core funding of the Max Planck Society (J.G.). The authors thank all members of the Toda and Guck lab for their support. We also thank Kirsten Hein, Oliver Vellage and Yuchang Lin for their continous support, without which this manuscript would not be possible.

## AUTHOR CONTRIBUTIONS

Conceptualization, Hein, K., Toda, T., Möckel, C., Romero-Limon, H., Zaccone, A., Guck, J.; methodology, Hein, K., Toda, T., Möckel, C., Romero-Limon, H., Zaccone, A; investigation, Hein, K., Möckel, C., Romero-Limon, H., Karasinsky, A.; writing-–original draft, Hein, K., Romero-Limon, H.; writing-–review & editing, Hein, K., Toda, T., Möckel, C., Romero-Limon, H., Zaccone, A, Kayser, J., Möllmert, S.; funding acquisition, Toda, T., Guck, J.; resources, Toda, T., Guck, J.; supervision, Toda, T., Guck, J., Kayser, J., Möllmert, S.

## DECLARATION OF GENERATIVE AI AND AI-ASSISTED TECH-NOLOGIES

During the preparation of this work, the author(s) used Anthropics Claude Code and Open AI ChatGPT in order to aid with coding and analysis. Furthermore, Claude, DeepL, and Overleaf’s own AI implementation were used to help with writing and translation of the manuscript. After using these tools or services, the authors reviewed and edited the content as needed and take full responsibility for the content of the publication.

